# Limited effects of dehydration on object discrimination in the novel object recognition paradigm in young and middle-aged male and female rats

**DOI:** 10.1101/2023.06.28.546884

**Authors:** Jessica Santollo, Ivanka L. Rainer, Lillian Swanz, Madelyn H. Steineker, Sahana Holla

## Abstract

Dehydration is associated with impaired cognitive function in humans. Limited animal research also suggests that disruptions in fluid homeostasis impair performance in cognitive tasks. We previously demonstrated that extracellular dehydration impaired performance in the novel object recognition memory test in a sex and gonadal hormone specific manner. The experiments in this report were designed to further characterize the behavioral effects of dehydration on cognitive function in male and female rats. In Experiment 1, we tested whether dehydration during the training trial in the novel object recognition paradigm would impact performance, while euhydrated, in the test trial. Regardless of hydration status during training, all groups spent more time investigating the novel object during the test trial. In Experiment 2, we tested whether aging exacerbated dehydration-induced impairments on test trial performance. Although aged animals spent less time investigating the objects and had reduced activity levels, all groups spent more time investigating the novel object, compared to the original object, during the test trial. Aged animals also had reduced water intake after water deprivation and, unlike the young adult rats, there was no sex difference in water intake. Together these results, in combination with our previous findings, suggest that disruptions in fluid homeostasis have limited effects on performance in the novel object recognition test and may only impact performance after specific types of fluid manipulations.

## Introduction

Multiple body systems are involved in defending body fluid homeostasis, maintenance of which is important because dehydration is associated with headaches, dizziness, fatigue, and in extreme cases, death. In addition, multiple reports in humans suggest that mild to moderate dehydration is associated with impaired cognitive function [1-5]. For example, in adult men exercise-induced or heat exposure-induced dehydration impaired response timing for perceptual tasks and impaired short-term memory recall [1]. While impairments in cognitive function have been reported in both men and women, sex differences in the degree of impairments have also been observed [6, 7]. The elderly are especially vulnerable to dehydration due, in part, to decrease thirst sensitivity [8-10], however, only limited research has focused on the effect of dehydration on cognitive performance in this population. Two studies have reported that poor hydration was associated with impaired declarative memory and slow psychomotor speed in health older adults (mean age 63 and 60 years, respectively) [11, 12]. While more research in particularly vulnerable populations is needed, there is sufficient evidence to demonstrate that dehydration negatively impacts cognitive function across a range of ages in humans.

Despite numerous studies in humans reporting negative effects of dehydration on cognitive function, there is limited work in laboratory animals characterizing this phenomenon. These types of studies are crucial for understanding the mechanisms by which low body fluids impact cognitive function. To our knowledge, there are only two reports in laboratory animals. In male mice, dehydration induced by 24 or 48 h water deprivation impaired memory recall in the Y-maze spatial memory test [13]. In this study, the mice were dehydrated during both the training and the testing trials, leaving open the question of whether dehydration impacts memory formation, recall, or both. We previously demonstrated in rats that dehydration during the test trial impaired performance (recall) in the novel object recognition test as a function of sex and gonadal hormone status [14]. Males, diestrous females, and ovariectomized females were not able to discriminate between the novel and original object when dehydrated, but estrous females were not affected by dehydration. Furthermore, we demonstrated that the impaired memory test performance occurred after extracellular dehydration (induced by treatment with the diuretic furosemide) but not intracellular dehydration (induced by treatment with hypertonic saline).

The goals for these studies were to further characterize dehydration-induced impairments in cognitive function in rats through use of the novel object recognition (NOR) test. We previously demonstrated that extracellular dehydration during testing impaired performance in the NOR in a sex specific manner [14]. Our first experiment in this study, therefore, tested the hypothesis that dehydration during training impairs subsequent euhydrated performance in the test (recall) trial in a sex and estrous cycle phase specific manner. Next, we tested the hypothesis that aging exacerbates dehydration-induced memory impairments in the novel object recognition test, as our previous work was limited to young adult rats [14]. We used 24 h water deprivation instead of furosemide treatment in the present studies, as it is a more natural mechanism for inducing dehydration.

## Materials and Methods

### Animals and Housing

Young adult (2–3-month-old) and middle aged (MA; retired breeders, 10–12-month-old) male and female Sprague Dawley rats (Envigo) were used. Animals were housed on a 12:12 h light:dark cycle with lights on at 700 h. All animals had *ad libitum* access to rodent chow (Teklad 2018) and tap water unless otherwise indicated. Animals were pair housed in standard shoebox cages until three days before the protocol began, at which point they were singly housed. During the early part of the light cycle, body weights were recorded and female rats in Experiment 1 had vaginal cytology samples collected for two weeks prior to testing, as previously described [15]. Estrous stage corresponded to the previous 12 h dark phase and the following 12 h light phase [16]. All experimental protocols were approved by the Animal Care and Use Committee at the University of Kentucky, and handling and care of the rats was in accordance with the *National Institutes of Health Guide for Care and Use of Laboratory Animals*.

### Open Field and Objects

All training and testing occurred in an open field measuring 75 cm wide X 75 cm long X 45 cm high. Grid lines on the open field divided it into 16 squares which were used to assess activity. The objects used for Experiment 1 were a clear plastic cylindrical jar and a martini glass. Both objects were similar in height and width (measured from the top of the martini glass). The objects used in Experiment 2 were a green bulb shaped spray bottle and a blue cylindrical spray bottle. The spray bottles were similar in height but differed in shape, color, and texture. Previous pilot studies in our laboratory found no differences in exploratory behavior between these pairs of objects [14].

### NOR Scoring

All sessions were video recorded for subsequent scoring of object exploration and activity. Object exploration was quantified in seconds (s) as time spent sniffing, licking, or touching the objects with the rat’s head or front paws during the trial [14]. If the rat’s rear or sides touched the object, this interaction was not included in the exploration time. Activity was quantified as the number of grid lines crossed during the trial. A line was counted as crossed when both front paws passed over the grid line. Videos were manually scored by at least two individuals blinded to the experimental conditions with an inter-scorer reliability of at least 90%.

### Experiment 1. Does dehydration during training impair performance during novel object recognition testing?

All rats (n = 48; 16 males, 32 females) were habituated to the empty open-field box for a 10 min period on Day 1. After habituation, rats were returned to their home cages and half of the subjects had their water bottles removed (n = 24). The following day (Day 2) consisted of a 3-min training trial during which rats were exposed to two identical objects in the open field box. Rats were then returned to their home cages and given pre-weighted water bottles. Four hours later, water bottles were weighed, and rats were placed back into the open field for the test trial. During the 3-min testing trial, rats were exposed to one of the original objects and a novel object. Habituation and training occurred during the first half of the light cycle (0900-1100). Testing occurred in the second half of the light cycle (1300-1500). This one day protocol was based on previous research [17]. We used this one day protocol instead of our multi-day protocol (Experiment 2 and [14]) to ensure the females were exposed to the fluid manipulations during the same stage of the estrous cycle (water deprived during the previous dark phase) as the training trial (trial conducted in the early light phase after dark phase water deprivation). Females were grouped into either a low estradiol group (n = 16), diestrus 1/diestrus 2, or a high estradiol group (n = 16), proestrus or estrus.

### Experiment 2. Does age influence dehydration-induced impairments in object recognition?

To test the effect of aging on dehydration-induced impairments in object recognition, we used our previously established 5 day protocol [14]. On Day 1, all rats (n = 104; 53 males, 51 females, 48 young adult, 56 MA) were habituated to the open field for 10 min. Training (Days 2-4), consisted of exposure to two identical objects in the open field for 5 min. After the training session on Day 4, half of the rats (n = 51) had their water bottles removed for 24 h. On Day 5, the testing trial occurred which consisted of exposure to one original object and one novel object for 5 min. After testing, all rats were returned to their home cages with pre-weighted water bottles and intake was measured after 30 min. All habituation, training, and testing occurred during the first half of the light cycle (0900-1100). Because the MA females did not have consistent 4–5-day estrous cycles, we did not group the females into different cycle stages as in Experiment 1.

### Data Analysis

Data are presented as means ± SEM throughout. All data were analyzed using the software packages Statistica. The discrimination index was calculated as (time spent with novel-time spent with original)/(time spent with novel + time spent with original). In Experiment 1, object exploration time was analyzed with a mixed factor ANOVA with Group and Hydration as between subject variables and Object (original and novel) as within subject variables. The discrimination index, activity levels, and water intake were analyzed with a two factor ANOVA (group by hydration). In Experiment 2, object exploration time was analyzed with a mixed factor ANOVA with Sex, Hydration, and Age as between subject variables and Object (original and novel) as within subject variables. The discrimination index, activity levels, and water intake were analyzed with a three factor ANOVA (sex by hydration by age). Newman Keuls post hoc tests were used to follow up significant ANOVA results.

## Results

### Experiment 1. Does dehydration during training impair performance during novel object recognition testing?

Although we previously reported that dehydration during novel object recognition testing impaired performance as a function of sex and stage of estrous cycle [14], it is unknown if dehydration during the training phase influences euhydrated performance in the recognition trial. Rats were, therefore, dehydrated or hydrated during the training trial and tested in a hydrated state. Time spend exploring the objects was influenced by the object, F_(1,42)_ = 30.71, p < 0.0001, but there was no effect of group, F_(2,42)_ = 1.68, p = ns, hydration, F_(1,42)_ = 1.34, p = ns, or any two or three way interactions (Figure 1A). Regardless of group or hydration state, rats spent more time exploring the novel, compared to the original, object, p < 0.05. The discrimination index was not influenced by group, F_(2,42)_= 1.63, p = ns, hydration, F_(1,42)_= 0.02, p = ns, or an interaction between group and hydration, F_(2,42)_ = 0.20, p < ns (Figure 1B).

**Figure 1.**
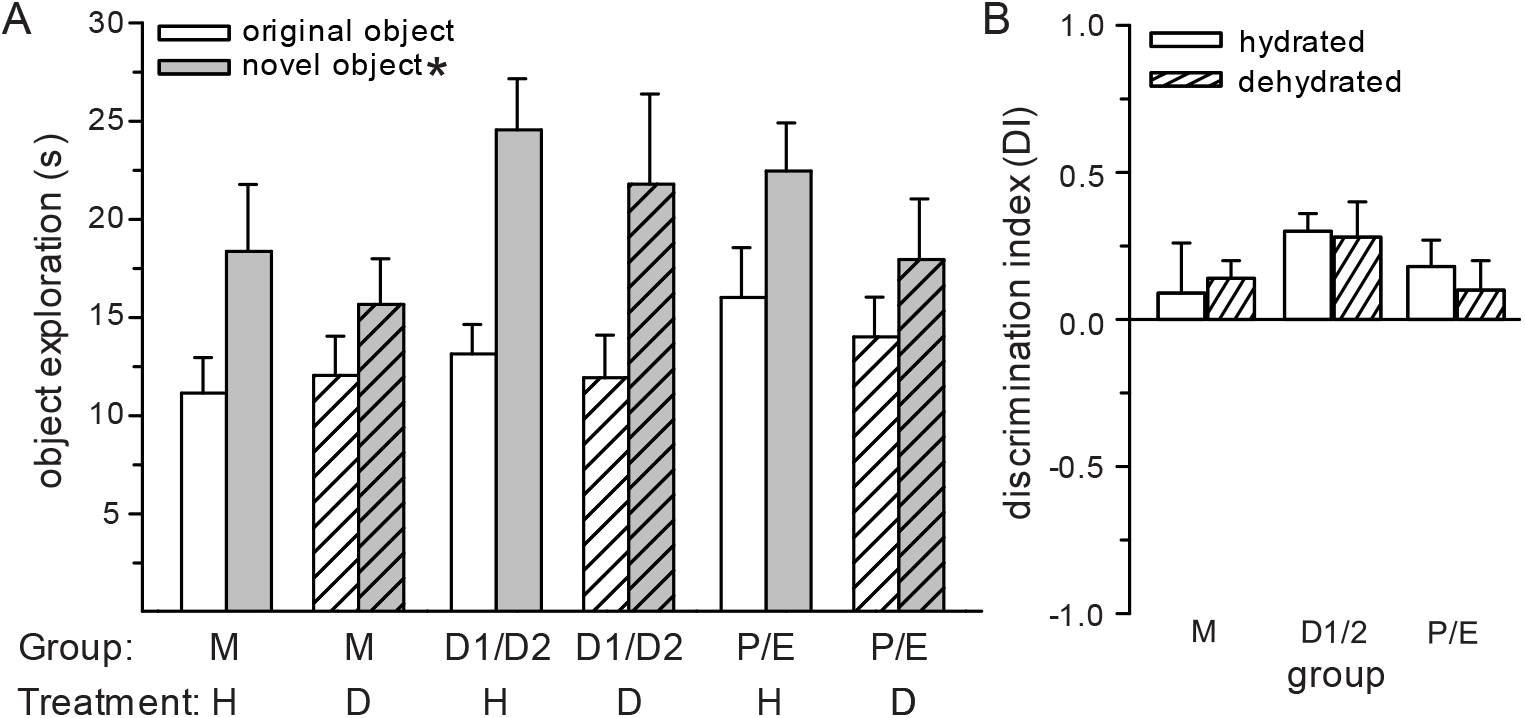
Dehydration during training had no effect on euhydrated performance in the novel object recognition test. (A) Regardless of group or hydration status, time spent investigating the novel object was significantly greater than time spent investigating the original object. (B) There were no differences in the discrimination index between any group. *Greater than original object, p < 0.05.

The number of lines crossed in the open field was analyzed as a proxy for activity levels to ensure that inactivity did not confound exploration of the objects. The number of lines crossed was influenced by group, F_(2,42)_= 7.83, p < 0.001, but not hydration, F_= 0.06_, p = ns, or an interaction between group and hydration, F_(2,42)_ = 0.06, p = ns (Figure 2A). Post hoc tests demonstrated that both the D1/D2 and P/E female groups crossed more grid lines than males, p < 0.05. Dehydration was confirmed by measuring water intake after the training trial. Water intake was influenced by hydration, F_(1,42)_= 113.9, p < 0.0001, but not group, F_(2,42)_= 2.59, p = 0.08, or a group by hydration interaction, F_(2,42)_= 1.58, p = ns (Figure 2B). Regardless of group, dehydrated rats consumed more water than hydrated rats, p < 0.05.

**Figure 2.**
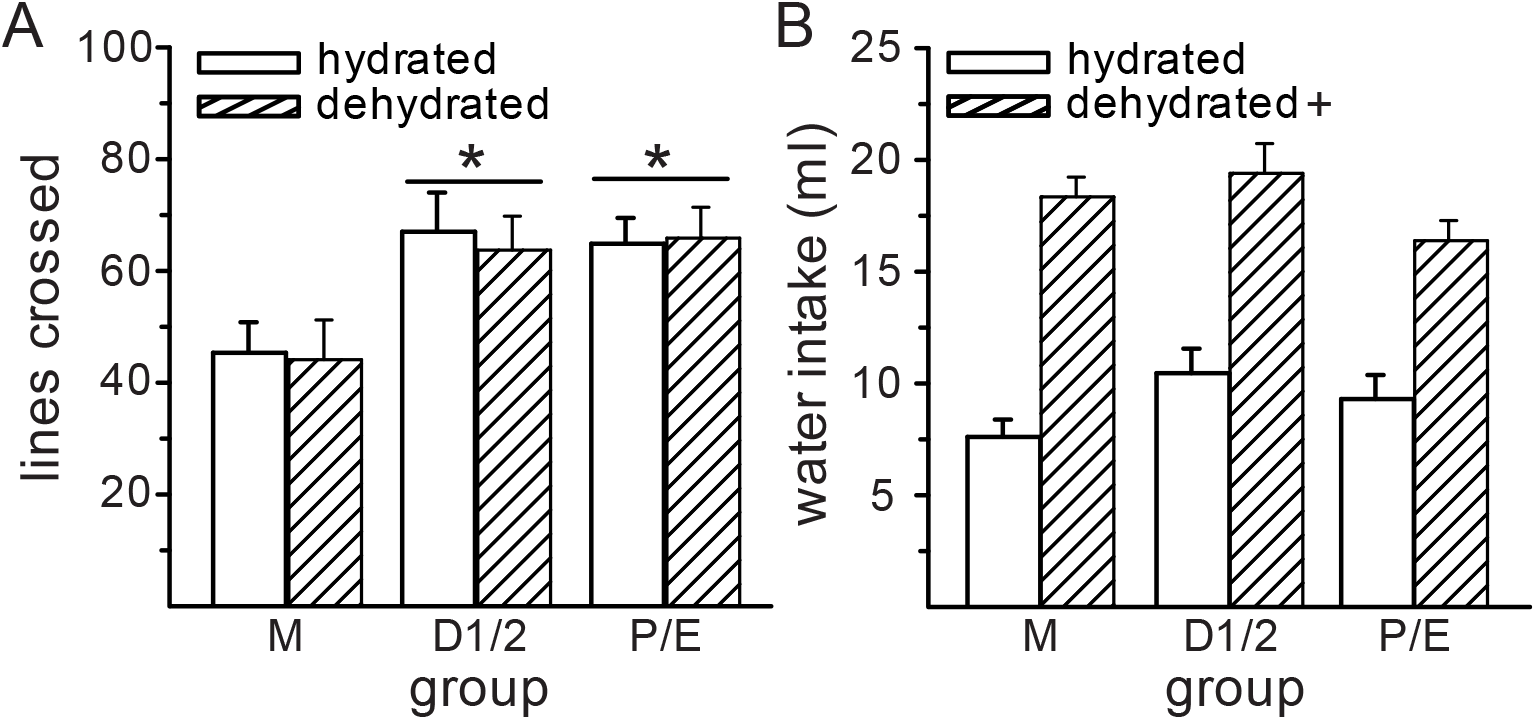
Activity and water intake measures. (A) Both groups of females were more active, as indicated by the number of lines crossed in the open field, compared to males. Dehydration during the training phase had no effect on activity levels during the test phase. (B) Dehydrated rats consumed more water in the 4 h period between training and testing that hydrated rats. *Greater than males, p < 0.05. ^+^Greater than hydrated rats, p < 0.05.

### Experiment 2. Does age influence dehydration-induced impairments in object recognition?

Although we previously reported that dehydration during novel object recognition testing impaired performance as a function of sex and stage of estrous cycle, it is unclear if aging exacerbates performance impairments. This experiment, therefore, compared dehydration-induced impairments in the NOR test between young adult and middle-aged rats. Time spend exploring the objects was influenced by the object, F_(1,96)_= 16.2, p < 0.001, age, F_(1,96)_= 92.59, p < 0.0001, and an interaction between age and sex, F_(1,96)_= 9.78, p < 0.01 (Figure 3A). Regardless of age, sex, or hydration status, all groups spent more time exploring the novel, compared to the original, object, p < 0.05. While young male and female rats spent a similar amount of time exploring the objects, MA females spent less time exploring the objects than the young rats, and MA males spent less time exploring the objects than either the MA females or the young rats, p < 0.05. There was no effect of hydration, F_(1,96)_= 0.16, p = ns, or any additional two or three way interactions. The discrimination index was not influenced by age, F_(1,96)_= 0.04, p = ns, sex, F_(1,96)_= 0.70, p = ns, hydration, F_(1,96)_= 0.31, p = ns, or any two or three way interactions (Figure 3B).

**Figure 3.**
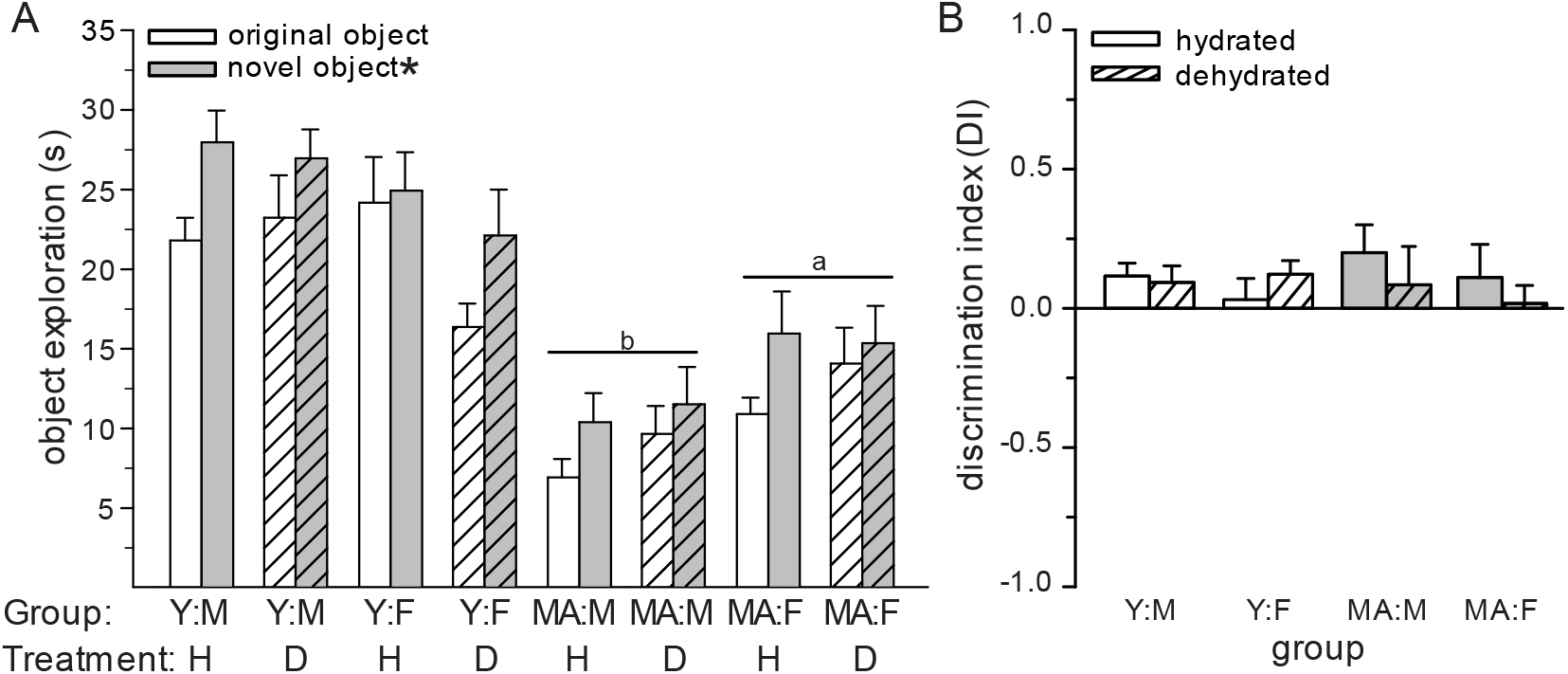
Dehydration and age had no effect on performance in the novel object recognition test. (A) Regardless of age, sex, or hydration status, time spent investigating the novel object was significantly greater than time spent investigating the original object. Object exploration time was reduced in the MA females, compared the young male and young female groups. Object exploration time in the MA males was less than all other groups. (B) There were no differences in the discrimination index between any group. *Greater than original object, p < 0.05. ^a^Less than young males and young females, p < 0.05. ^b^Less than young males, young females, and MA females, p < 0.05.

To ensure that inactivity did not impair performance during the test trial, the number of lines crossed in the open field was analyzed as a proxy for activity levels. The number of lines crossed was influenced by age, F_(1,96)_= 119.2, p < 0.0001, sex, F_(1,96)_= 25.74, p < 0.0001, an interaction between age and sex, F_(1,96)_= 5.51, p < 0.05, and an interaction between age, sex, and hydration, F_(1,96)_= 8.79, p < 0.001 (Figure 4A). Young rats crossed more lines than the MA rats, p < 0.05. Females crossed more lines than males, p < 0.05. Young females crossed more lines than all other groups. Young males crossed more lines than both male and female MA rats and female MA rats crossed more lines than male MA rats, p <0.05. Although there was a significant three-way interaction, no additionally meaningful group differences were detected. Dehydration was confirmed by measuring water intake after the test trial. Water intake was influenced by hydration, F_(1,96)_= 232.02, p < 0.0001, age, F_(1,96)_= 10.82, p < 0.01, and an interaction between age, sex, and hydration, F_(1,96)_= 4.92, p < 0.05 (Figure 4B). As expected, dehydrated rats consumed more water than hydrated rats, p < 0.05. Young rats consumed more water compared to MA rats, p < 0.05. Dehydration induced water intake was greater in young male, compared to young female, rats, p < 0.05. This sex difference was not detected in the MA rats.

**Figure 4.**
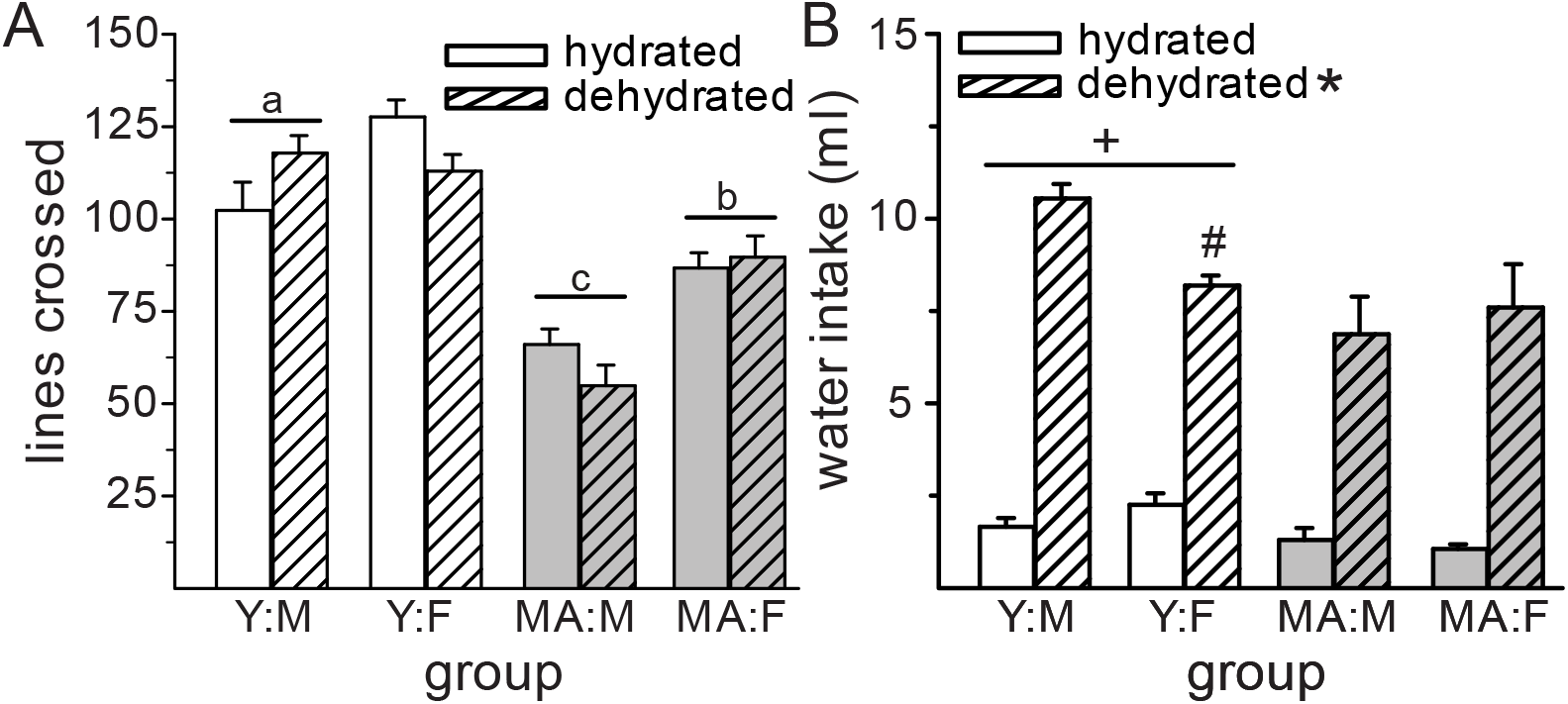
Activity and water intake measures. (A) Young females were more active, as indicated by the number of lines crossed in the open field, compared to all other groups. Young males were more active than the MA males and females. MA females were more active than the MA males. Dehydration had no effect on activity levels during the test phase. (B) Dehydrated rats consumed more water after the test trial than hydrated rats. Young rats consumed more water than MA rats. In young rats, dehydrated males consumed more water than dehydrated females, but this sex difference was not detected in MA rats. ^a^Less than young females, p < 0.05. ^b^Less than young females and males, p < 0.05. ^c^Less than all other groups, p < 0.05. *Greater than hydrated rats, p < 0.05. +Greater than MA rats, < 0.05. ^#^Less than dehydrated young male rats, p < 0.05.

## Discussion

The goal of this series of experiments was to further characterize the behavioral effects of dehydration on cognitive function in a rodent model. As such, we examined the effect of 24 h water deprivation in male and female rats on performance in the novel object recognition paradigm. In Experiment 1, we found that dehydration during the training trial had no effect on object discrimination in the test trial. In Experiment 2, we found that dehydration during the test trial also had no effect on object discrimination in either young adult or middle-aged rats. Age,however, was associated with reduced activity levels and reduced deprivation-induced water intake. Despite a few reports of impaired performance in memory trials as a function of dehydration in rodents [13, 14], the present data suggest that either dehydration has very limited impairments in the NOR paradigm or that subtleties in the fluid manipulations may have large impacts on performance.

We have previously demonstrated that extracellular dehydration during NOR testing impaired object discrimination in males, diestrous females, and ovariectomized females. We, therefore, tested whether dehydration during NOR training would impair object discrimination when rats were tested in a hydrated state. There was, however, no effect of dehydration during training on test performance. All groups, regardless of sex/estrous cycle stage or hydration status, spent more time investigating the novel object, compared to the original object. This suggests that either dehydration does not impair training or that our paradigm was not able to detect an effect of dehydration. Our paradigm could have been insensitive to the dehydration manipulation for several reasons. First, task difficulty can affect performance in the NOR [18]. It is possible that the objects used in this experiment (cylindrical jar and a martini glass) were too easy to discriminate. The objects used in Experiment 2 were the same objects that we observed dehydration induced impairments with in our previous report [14], therefore, a follow up study should repeat this experiment with those objects to rule out task difficulty as a factor. It is also possible that 24 h water deprivation is not as severe as treatment with furosemide, which was the mechanism used to induce dehydration in our previous report [14]. While it is possible that 24 h water deprivation is not severe enough to negatively affect the training trial, the report by Faraco and colleagues found that 24 h water deprivation was sufficient to impair performance [13]. There are, however, important differences between these two studies including different rodent species (mice vs rats), different paradigms (Y-maze vs NOR), and different aspects of the paradigm subjected to dehydration (training + testing vs training). Regardless of these possibilities, dehydration during the NOR training does not have a robust, if any, effect on discrimination during the test trial.

Experiment 2 was designed to determine if aging exacerbated dehydration-induced impairments in NOR performance. Our testing paradigm (5 day protocol) was identical to our previous report, where we observed that dehydration impaired discrimination of the novel and original object in males, diestrous female, and ovariectomized females [14]. We found, however, that dehydration had no effect in any group. All groups regardless of age, sex, or hydration spent more time investigating the novel object, compared to the original object. This was surprising as we failed to replicate our previously reported impairments in test performance. The only difference between this study and our previous report was that we used 24 h water deprivation to induced dehydration here, but in the previous study we used furosemide treatment [14]. Again, it is possible that the severity of dehydration induced by 24 h water deprivation is not as severe as furosemide treatment. It is also possible that due to the multiple variables, our experiment was not sufficiently powered to detect differences. All groups, however, had a positive discrimination index (indicating more time spent with the novel object) and separate ANOVAs of the young and MA groups and males and females did not detect any effects of dehydration. It was also surprising that dehydration had no effect on test performance in the MA group as we predicted this group would be more sensitive to impairments given that cognition declines as a function of age [19-21]. It is likely that the age group used here (10-12 months old) was not old enough to have significantly impaired memory function. Indeed, age related impairments in place discrimination in the Morris Water Maze have been reported to be significantly impaired in 17 and 24 month old rats, but only mildly impaired in 11 month old rats and impairments in working memory were only observed at 24 months of age [19]. Future studies should use older rats to determine if dehydration-induced impairments in memory are exacerbated by aging.

Sex and age impacted activity levels during the test trials. In Experiment 1, regardless of hydration status, both groups of female rats were more active that the males in the test trial. This is not surprising given the well documented sex difference in activity in rodents [22]. In Experiment 2, effects of sex and age impacted activity levels regardless of hydration. Females were more active than their age-matched male counterparts. In addition, younger rats were more active than the MA rats. While this was not surprising, as it has been previously reported that aging is associated with decreased activity levels [23, 24], there are conflicting reports of the sex difference in locomotor activity persisting in aged rats [25, 26]. These findings strengthen previous findings that aged female rats are more active than aged male rats [26]. The aged animals also had reduced interaction times with the objects during the test trial, which is likely a result of their overall reduced activity levels, especially since there was a parallel sex difference with aged females spending more time investigating the objects than aged males. Despite no effect of age on NOR performance, the age-related differences in locomotor activity at least demonstrate physiological differences between the two age groups and suggest that physiological differences (at least related to locomotor activity) are affect by aging earlier than any aging-related impairments in memory recall in the NOR paradigm.

Water intake was the only variable in these experiments that was significantly affected by 24 h water deprivation. As expected, dehydrated rats consumed more water compared to euhydrated rats. In Experiment 1, there were no differences in water intake between male, D1/D2 females, or P/E females. There were, however, sex differences in dehydration-induced water intake in Experiment 2. Young males consumed more water after 24 h water deprivation compared to young females, supporting previously reported sex differences [27, 28]. It is unclear why the sex difference was not detected in Experiment 1, but it could be due to the different time intervals of intake. In Experiment 1, water intake was measured after 4 h (just prior to the test trial), while in Experiment 2, water intake was measured after 30 min. Rehydration after water deprivation occurs rapidly, usually within 10-20 minutes, therefore the 4 h interval also captured diurnal *ad libitum* intake which could have masked the sex difference. Dehydrated MA rats consumed less water than dehydrated young rats and there was no sex difference in dehydration-induced water intake in the MA rats. To our knowledge there are only two reports examining water intake after 24 h water deprivation in aged male and female rats. In the first report, males consumed more water than females but there were no age-related effects in water intake [28]. It is, however, unclear if the sex difference in water intake was driven by the young rats, as there is no mention of whether an age by sex interaction was significant or not and according to the graphed data, water intake at 10 months of age was identical in males and females [28]. In the second report, no sex differences were reported, but young rats consumed more water than aged rats [29]. The data in this report, however, were expressed as ml/bw, which we have argued is inappropriate as body weight does not correlate with water intake in females and only correlates in males under certain conditions [15, 27]. Our findings here are therefore in agreement with the reported sex difference in young rats in [28] and the age-related decrease in water intake reported in [29]. We, therefore, conclude that the sex difference in water intake stimulated by 24 h water deprivation is absent in MA rats.

Although research in human subjects suggests that mulitple aspects of cognitive function are impacted by moderate, or even mild, dehydration, we failed to identify effects of dehydration on NOR performance in this series of experiments. Our previous research demonstrated that extracellular dehydration, induced by furosemide treatment, impaired performance in the NOR in males, diestrous females, and ovariectomized females [14]. Here, however, we failed to find an effect of 24 h water deprivation during the training trial on euhydrated performance in the test trial. We also failed to replicate our previous finding that dehydration during testing impaired object discrimination. The most likely explanation is that while dehydration attained by furosemide treatment can impact test performance, dehydration induced by 24 h water deprivation cannot. This was surprising as it has previously been reported in mice that 24 h water deprivation impacts spatial memory [13]. It will be important to rule out the confounds discussed above before concluding that dehydration only has minor effects of memory performance in the NOR test. In addition, different behavioral tests that probe other types of memory (for example more spatial memory tasks) should be conducted with dehydration manipulations. Given the well documented effects that dehydration has on cognition in humans, animal studies will be critical to understand how low fluid levels impact learning and memory and help guide prevention techniques and therapeutics. Furthermore, given rising global temperatures and dramatic changes in rainfall, wild animals are going to be more likely to encounter unpredictable access to water and as a result dehydration. Hence, dehydration-induced cognitive impairments could have extremely detrimental effects on wild animals attempting to survive in a rapidly changing climate [30]. As such, understanding the underlying neurobiology will be necessary in facilitating interventions.

## Acknowledgments and Support

We thank Andrea A. Edwards for her assistance with animal husbandry. This work was supported by NIH grant DA035150 and University of Kentucky, College of Arts and Sciences Start-Up Funds.

## Notes

### Competing Interest Statement

The authors have declared no competing interest.

